# Eliminating steam pops and improving lesion safety in atrial ablation with conductive hydrogels

**DOI:** 10.1101/2023.10.03.560762

**Authors:** Allison Post, Lukas Jaworski, Drew Bernard, Derek Bashe, Shang Gao, Abbey Nkansah, Haichong Zhang, Elizabeth Cosgriff-Hernandez, Mehdi Razavi

**Author notes:** Corresponding Author: Mehdi Razavi, MD;, 6770 Bertner Avenue, MC 2-255 Houston, TX 77030.

## Abstract

**Background:** Atrial fibrillation (AF) is a significant burden worldwide, and the existing treatments leave much to be desired. There are, however, opportunities to improve the safety and efficacy of the most popular treatment, radiofrequency (RF) cardiac ablation, using conductive hydrogels as an ablation mediator.

**Methods:** Lesions were created in ex vivo ventricular tissues using bare metal traditional RF catheters and three different hydrogels with varying conductivities to assess the effect of conductivity on lesion formation. Similar procedures were performed in atrial/esophageal tissue stacks to mimic physiological AF ablation and demonstrate the initial safety profile of conductive hydrogel-mediated ablation.

**Results:** The hydrogel mediated lesions were overall shallower and narrower than bare metal, and also exhibited less char and improved lesion homogeneity. The hydrogel also eliminated steam pops. Finally, the hydrogel appeared to be more thermally protective of the esophagus in the atrial/esophageal tissue stack, greatly reducing the lesion formation on the esophagus while still achieving transmural lesions in the atrial tissue.

**Conclusions:** Hydrogel-mediated RF ablation holds promise as a novel method to improve ablation outcomes for AF patients. Future work will confirm this in vivo and establish the chemistry required to create a conductive hydrogel coating for RF ablation catheters.

## INTRODUCTION

Atrial fibrillation (AF) affects up to 6 million people, with 750,000 hospitalizations per year costing $28 billion annually in the US alone^1^. If left untreated, AF can lead to stroke, heart failure, and other heart-related complications. Current treatment methods include medication and/or elimination of the arrhythmia-causing cells and circuits via cardiac ablation. Antiarrhythmic drug therapy alone is only effective in resetting rhythm in approximately 57% of patients ^2^, and under the right conditions, these drugs can actually be proarrhythmic ^3^. When medical management fails or is incompatible with the patient, then cardiac ablation is used. Cardiac ablation, while typically more effective than medical management, still suffers from recurrence rates of up to 45%^4^. This is due mostly to the tradeoff required in delivering durable lesions and protecting the surrounding, healthy tissue, resulting in the potential for re-entrant circuit reconnection ^5^.

The clinical gold standard for cardiac ablation is radiofrequency (RF) ablation. Although RF ablation can create stable lesions, delivery of ablation is limited by the poor thermal contact of the metal tip of RF catheters and the uneven surfaces of the myocardium, resulting in the potential for steam pop formation, tissue char, thrombus formation, and thermal damage to surrounding tissues such as nerves and the esophagus^6^. Alternatives have been explored and marketed to address some of the limitations of RF ablation, but still have not achieved markedly improved efficacy or safety. Once such technology, cryoablation, can reduce procedure times and lower the risk of pericardial effusion and tamponade, but it does not have an improved efficacy in maintaining sinus rhythm and actually increases the risk of phrenic nerve damage ^7^. Even more recently, pulsed field ablation (PFA) has shown some promise in reducing overall complications due to some tissue selectivity during ablation. Although initial data looks promising, more long-term data and randomized controlled trials are still needed for validation and widespread adoption ^8^.

Despite new technologies becoming available on the market, RF ablation remains the most utilized method for treating AF. However, there remains a need to thermally protect surrounding tissues and structures and reduce the incidence of steam pop during RF ablation procedures. Therefore, there is a clear and pressing need for a solution that can provide controlled delivery of radio-frequency ablation that could be easily integrated into current catheter and generator technology. In the work presented here, we explore a range of conductive hydrogels as a thermal mediator of RF ablation for more controlled, safer lesion formation using porcine tissue in an ex vivo ablation model.

## METHODS

Hydrogel synthesis and conductivity characterization: Poly(ether urethane diacrylamide) (PEUDAm) hydrogel slabs were fabricated at 20wt% in water with 0.1wt% Irgacure 2595 under UV cure. 1cm punches were taken from the slabs and dehydrated until time of experimentation. The gels were rehydrated the day before the study in saline solutions of varying concentrations to vary the overall conductivity of the gel. Hydrogel candidates for RF ablation mediation were based on conductivity measurements made with electro-chemical impedance spectroscopy. The saline solutions selected were 0%, 0.5%, and 1.5% saline in water, correlating to 0, 6, and 15 mS/cm, respectively.

Tissue sample preparation: Fresh porcine hearts were obtained (Animal Technologies, Tyler, TX) one day after sacrifice and excision. Tissue samples were excised from the hearts using a square cutting guide to partition consistent sections from the left ventricle, right ventricle and septum of the heart. The tissue samples were then placed into a custom 3D printed sample alignment cage (**Figure 1 B**), serving to center the ablation lesion and subsequent lesion analysis steps. Tissue samples were kept wrapped in PBS-soaked gauze when not being ablated to retain hydration and prevent tissue necrosis.

**Figure 1:**
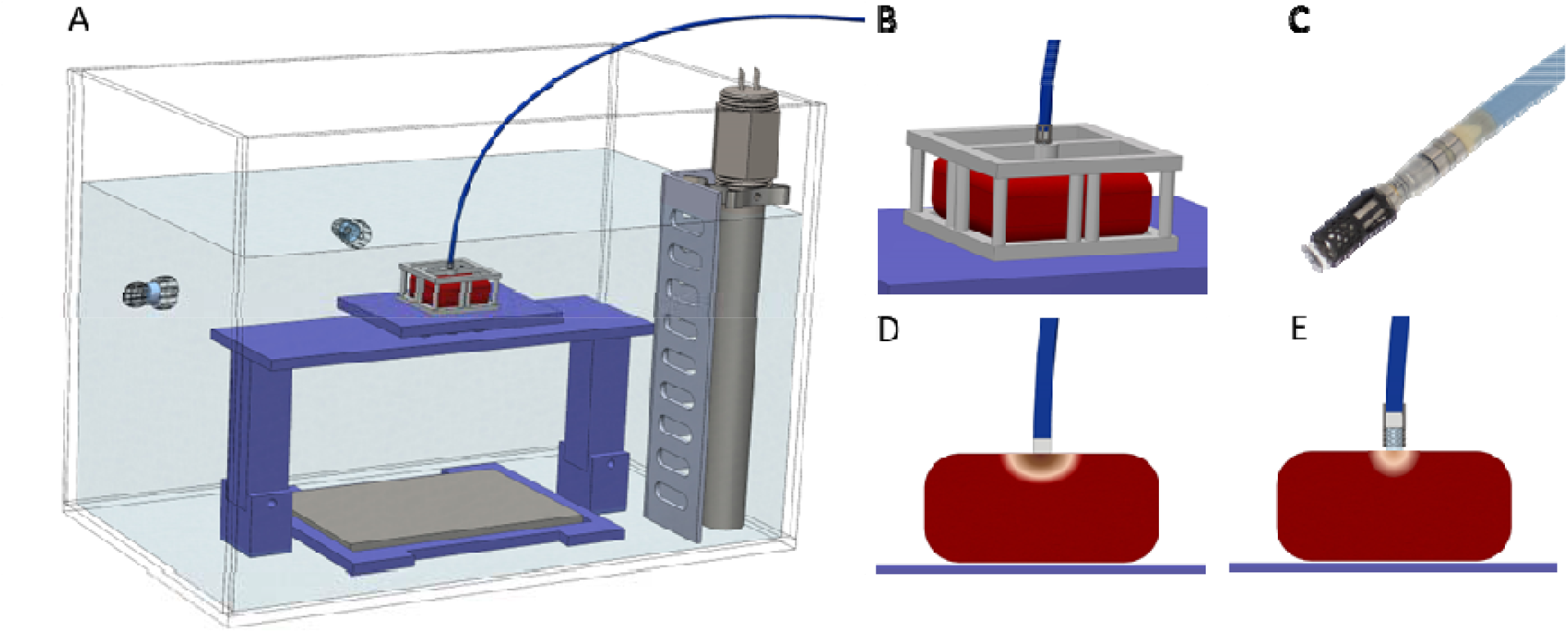
A. Full tank assembly for ex-vivo tissue ablation, including tissue stage, grounding pad, circulating water heater set to 37C, and 0.9% saline solution. B. Custom designed and printed tissue holder for ablation catheter alignment. C. Photo of custom designed and printed hydrogel punch holder fitted to the end of an RF ablation catheter. D. Figure of bare metal RF catheter ablating a tissue sample E. Figure of a hydrogel-mediated ablation with a modified RF catheter.

For assessment of thermal transfer of ablation, tissue stacks were created using porcine left atrial back wall and sections of the esophagus (Animal Technologies, Tyler, TX), as occurs naturally in human physiology. These stacks were placed on the tissue holder and held in place with bands for submersion in the ex vivo ablation tank (**Figure 2**). Ablations were focused on smooth, unstriated portions of the atria stacked on top of the esophageal tissue segment.

**Figure 2:**
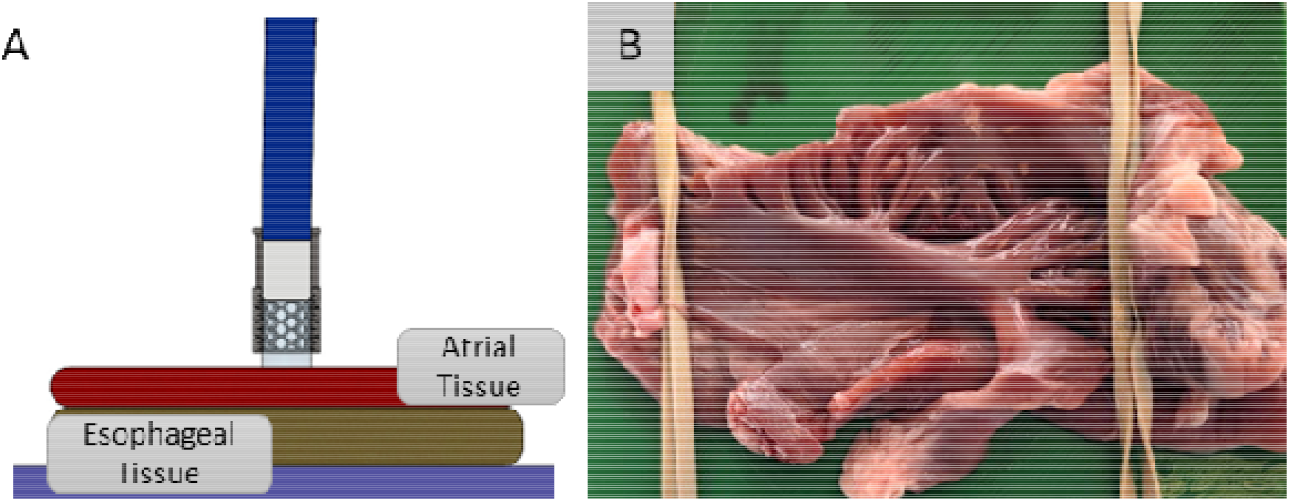
A. Figure of the atrial esophageal tissue stack ablation. B. Image of the stack on the tissue holder to be placed in the ex vivo ablation tank. Hydrogel punches and attachment to ablation catheter: Desiccated Poly(ether urethane diacrylamide) (PEUDAm) hydrogel discs were first rehydrated in either 0%, 0.5%, or 1.5% saline overnight. Cylindrical hydrogel sections from each concentration were prepared with a diameter equal to that of the ablation catheter tip (8Fr). The hydrogel samples were fixed onto the end of an irrigated 8-Fr ThermoCool™ SF ablation catheter (Biosense-Webster, Irvine CA) with a custom 3D printed catheter attachment (**Figure 1C**). The catheter attachment allowed the hydrogel to protrude from it slightly (.5 – 1 mm), allowing unimpeded contact with tissue (**Figure 1 C**).

Ablation parameters: The catheter tip, either bare metal or fitted with a hydrogel tip, was then centered in the tissue alignment cage prior to ablation (**Figure 1 D&E**). For tissue stacks, the ablator was held in place by the operator. The ablations were carried out inside of a 37° C temperature controlled saline tank using (Carto generator, Biosense Webster, Irvine CA), with a stainless-steel plate serving as the indifferent electrode, located approximately 6 cm below the tissue stage (**Figure 1 A**). Each sample underwent ablation at 30 watts for 30 seconds with a contact force of 20 grams. Steam pop occurrence was observed by the operator and recorded for each test. After ablation, samples were re-wrapped in PBS-soaked gauze and stored on ice overnight.

Tissue sample staining and imaging: Ablated tissue samples were removed from iced storage and sliced vertically through the center of the ablation lesion. 2,3,5-Triphenyltetrazolium chloride (TTC) powder was dissolved at 2% w/v in 0.9% saline solution and warmed to 37°C. Sliced samples were incubated for 20 minutes in the TTC solution to enhance the margins between the ablated tissue and the healthy tissue. Images of the stained ablated tissue were acquired after removal from the staining solution. Image analysis: Lesion depth and width were measured from the acquired images using ImageJ. Statistical analysis was performed using R ANOVA tool kit, with alpha = 0.05.

## RESULTS

Steam pop formation during ablation: Steam pops were observed only in the bare metal catheter ablation, occurring in 2 out of 6 burn events, or 33.3% (**Figure 3**). No steam pops were observed in any of the hydrogel mediated ablations.

**Figure 3:**
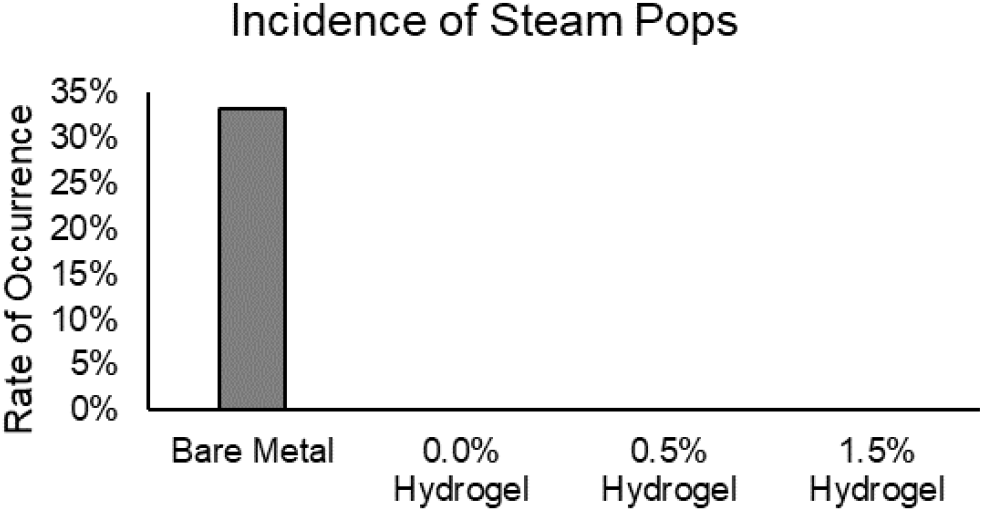
Incidence of steam pops during RF ablation of ventricular tissue determined by percentage of burns experienced a steam pop event (n=6).

### Lesion comparisons in ventricular tissue

Overall, burn depths and widths were significantly lower in the hydrogel mediated tissue than in the bare metal (**Figure 4 B, C**). There are significantly lower levels of char as well in the hydrogel mediated lesions compared with the bare metal lesions (**Figure 4 D, E**), and char width also varies significantly between hydrogel formulations (**Figure 4 D**).

**Figure 4:**
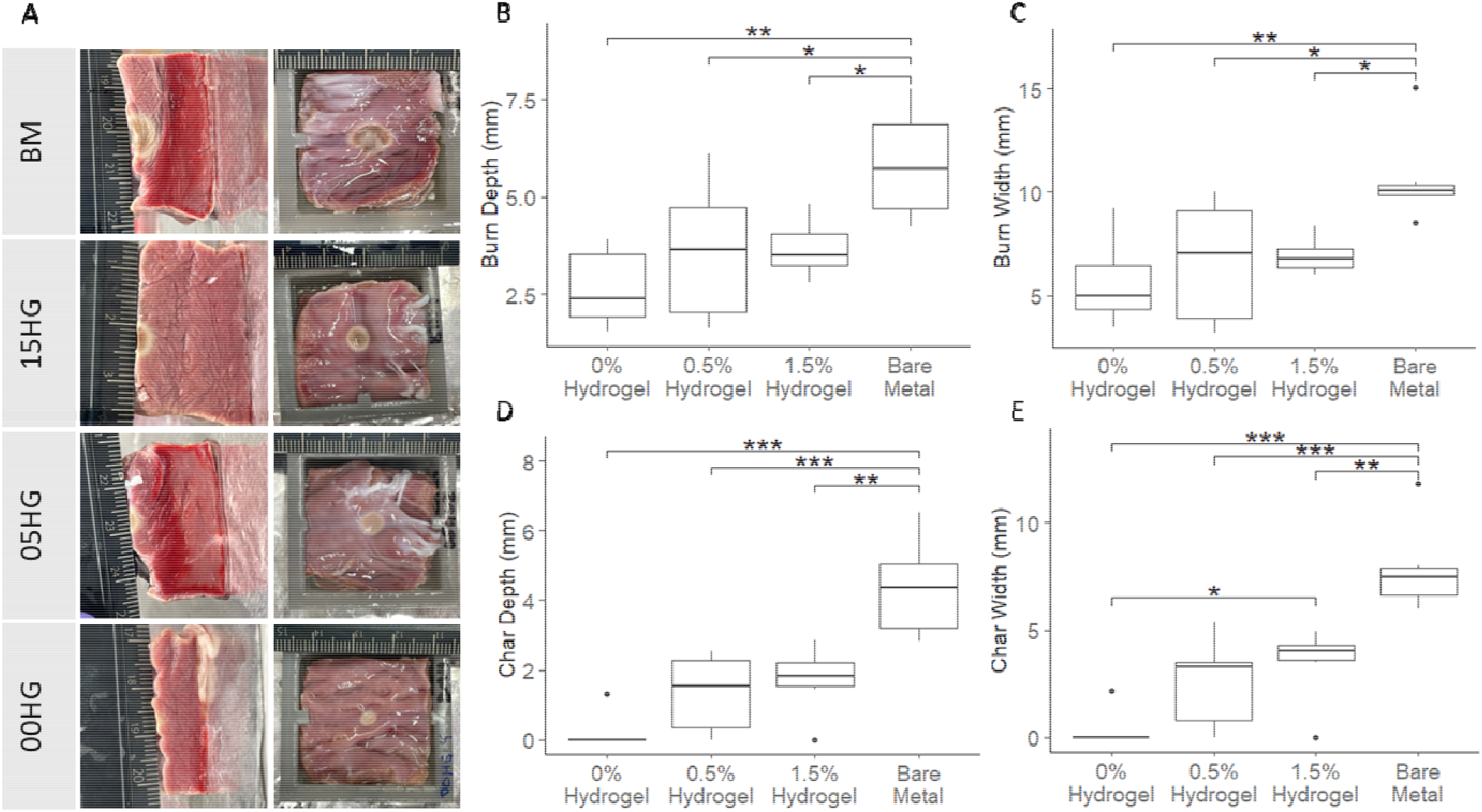
A. Representative images of cross sections of lesions from each test group. B. Measured lesion depth from cross-sectional images. C. Measured lesion width from ablated tissue images. D. Measured lesion char depth from cross-sectional images. E. Measured lesion char width from cross-sectional images.

### Lesion comparisons in atrial/esophageal tissue stacks

In the stacked tissue, the overall burn depth follows a similar trend to those seen in the thicker ventricular tissue, namely that bare metal has a significantly deeper lesion measurement (**Figure 5 B**). While not significant, the esophageal lesion depth and width appear to trend lower with a hydrogel mediated burn than with bare metal (**Figure 5 D, E**). All atrial burns were transmural (**Figure 5 D**).

**Figure 5:**
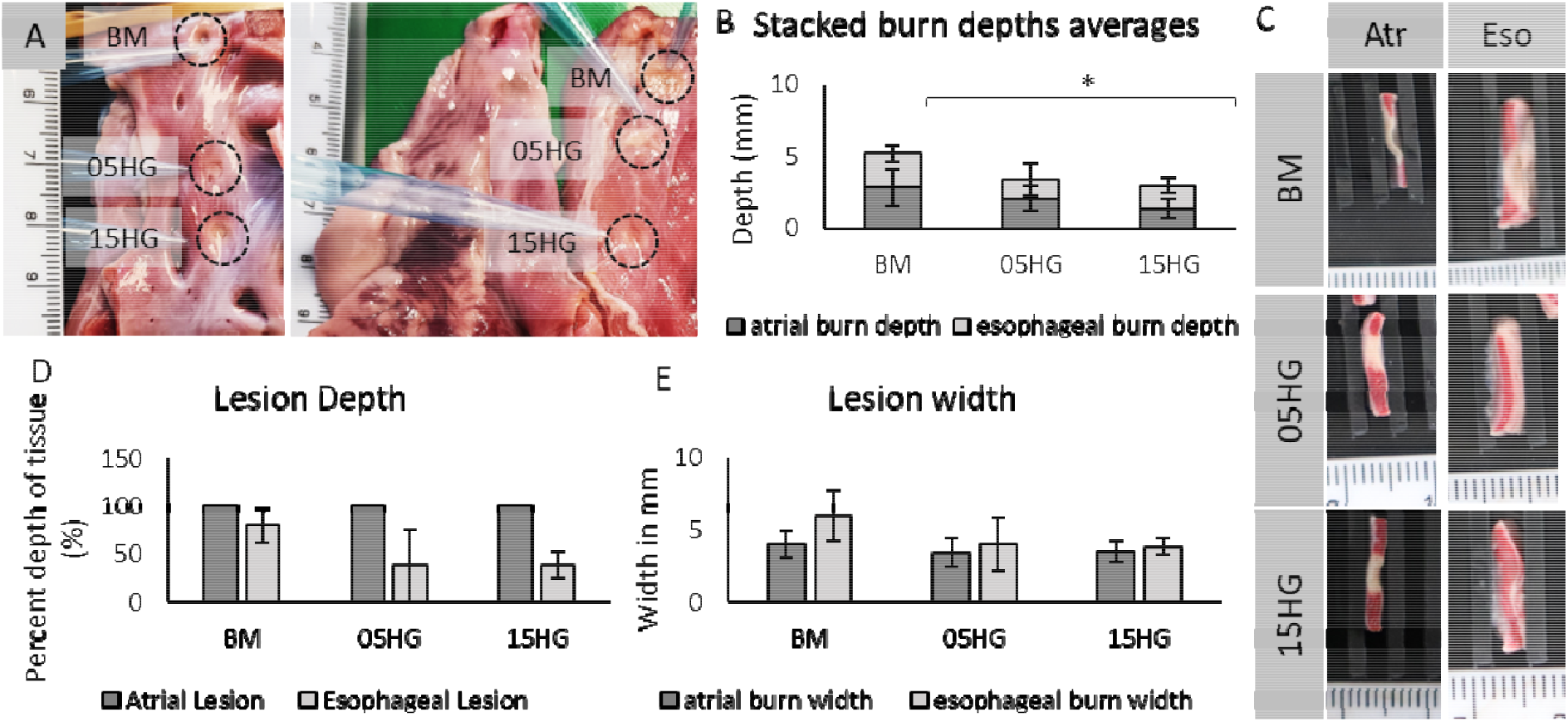
A. Representative images of ablation lesions on atrial endocardium (left) and esophagus epicardium (right), with lesions formed by bare metal (top), 0.5% saline hydrogel (middle), and 1.5% saline hydrogel (bottom) B. Average depth of the lesion in the tissue stacked for the bare metal (BM), 0.5% saline hydrogel (05HG), and 1.5% hydrogel (15HG) lesions. C. Representative cross-sectional areas of lesions in both atrial and esophageal tissue D. Measured average lesion depth by percent of tissue burned. E. Measured lesion width in mm.

**Figure 5:**
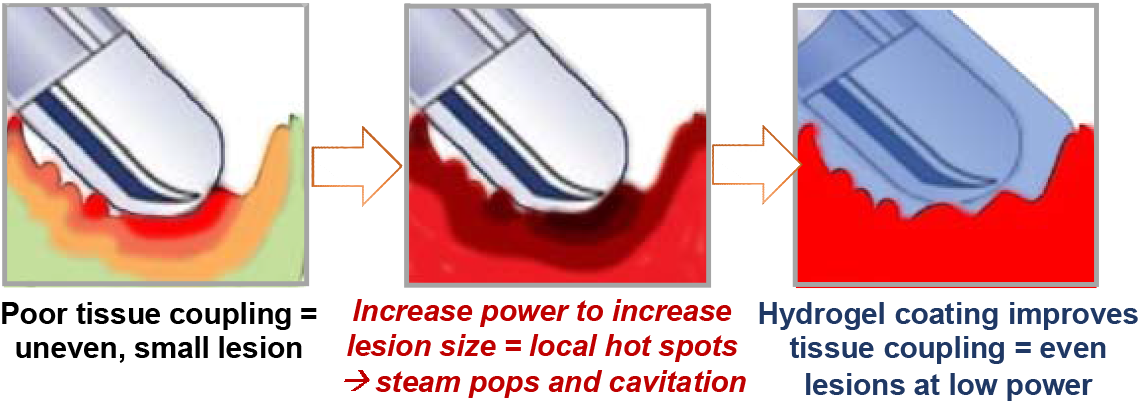
The underlying issue of contact, which ultimately affects lesion formation and steam pop formation, and the proposal of mechanism of improvement in steam pop reduction and homogeneous lesion formation by a mediating conductive hydrogel.

## DISCUSSION

The markedly absent steam pop observation in all hydrogel-mediated ablations is a vast improvement over the higher rate of steam pop formation (33%) in traditional RF ablation catheters. Even in an ex vivo experiment, this is an exciting development, because of the potential for clinical translation. The risk of steam pop formation remains one of the most common and yet potentially complicating events that can occur during routine catheter ablation. Current clinical data suggests mitigating these will lead to improved outcomes [REF].

The apparent elimination of steam pop formation is a non-trivial and likely complex combination of factors. The proposed hypothesis is that the soft, moldable hydrogel interface likely provides more even and consistent contact between the ablation catheter and the uneven tissue surface (Figure 5), providing better thermal regulation and transmission and potentially bubble trapping. Further investigation is required to understand the underpinning physics of this phenomenon.

In assessing the lesions in thick, ventricular tissue, we are able to understand some of the basic effects of the hydrogel mediated RF ablation compared to the traditional bare metal RF catheter. Namely, the hydrogel mediated lesions are smaller and more even than the bare metal lesions. The reduced measurements of char in the hydrogel mediated lesions implies a more even lesion with an improved safety profile. Namely, this lower area of char implies there is less microvascular damage, and therefore a lower risk of thrombus formation and lower risk for thromboembolic events.

From the data shown in **Figure 4**, it is obvious that the level of conductivity of the hydrogel is key in shaping the resulting lesion. As conductivity is increased from 0% to 0.5% and 1.5% saline, there appears to be a correlative increase in ablation lesion size. The correlation between hydrogel conductivity and resulting lesion characteristics provides an interesting opportunity for fine-tuning delivery of RF ablation for various applications. Lesion depths and widths could be modulated purely by a conductive hydrogel coating, with the added benefit of eliminating steam pops and risks for microvascular damage that results in a cascade of undesirable events.

More work will be required to understand the relationship between conductivity and resulting lesion size in a more granular manner to optimize a potential hydrogel coating for a traditional RF ablation catheter. Confirmation of this technology’s benefits in an in vivo setting will also be critical.

In assessing the lesions in atrial esophageal tissue stacks, we can begin to understand the clinical relevance of the hydrogel mediated RF ablation. In the data presented, all lesions regardless of RF mediator are transmural in the atrial tissue as would be desired in an ablation procedure, and all groups had similar lesion widths in the atrial tissue. The significant difference is the overall lesion depth, with hydrogel mediated RF ablation showing significantly less damage to the esophagus compared to bare metal. The protective nature of the hydrogel are then two-fold: ensures a transmural lesion without steam pop while thermally protecting the esophagus.

## CONCLUSION

To address the need for transmural atrial lesions with esophageal protection while also reducing steam pops, it appears that conductive hydrogels hold promise. The initial ex vivo data indicates safer lesion formation in an atrial setting and elimination of steam pops. Further in vivo data will be critical in confirming these findings and pushing this technology towards a viable catheter coating chemistry.

## ACKNOWLEDGEMENTS

Dr. Deborah Vela, THI Pathology for experimental design input; Houston Biosense Webster Clinical Team for lending RF ablation generator

## SOURCES OF FUNDING

This research was funded by the NIH National Heart, Lung, and Blood Institute (1R01HL162741), the Cizick Foundation, and the Roderick MacDonald Fund.

## Notes

### Competing Interest Statement

The authors have declared no competing interest.

### Summary of Updates

author list has been updated and affiliations have been updated.

